# Network Pharmacology Analysis of the active components and anticancer targets of Rhubarb

**DOI:** 10.1101/2021.01.28.428583

**Authors:** Hu Junrui, Duan Yongqiang, Cui Gongning, Luo Qiang, Xi Shanshan, Huang Rui, Ma Jun, Bai Min, Wu Hongyan

**Affiliations:** College of Basic Medicine, Gansu University of Chinese Medicine,Lanzhou 730000,China; Department of traditional Chinese Medicine,Yinchuan third people’s Hospital,Yinchuan 750001,China; Department of diabetes, Hengshui Hospital of traditional Chinese Medicine,Hengshui 053000,China; Department of traditional Chinese Medicine, Shenzhen Hospital of Shanghai University of traditional Chinese Medicine,Shenzhen 518001,China

## Abstract

To investigate the mechanisms and active components governing the anticancer activity of rhubarb.The TCMSP database was screened to identify the active components of rhubarb and Swiss target predictions were generated to predict their cellular targets. TTD and OMIM databases were used to predict tumor-related target genes. "Cytoscape" was used to construct drug targets. PPI network analysis, GO enrichment analysis and KEGG pathway analysis of the key targets were investigated using String and David databases. A total of 33 components and 116 corresponding targets were screened. Amongst them, the key active compounds in rhubarb included emodin, aloe emodin, β-sitosterol, emodin methyl ether and rhein, which were predicted to target TP53, AKT1, STAT3, PIK3CA, HRAS, and VEGFA. GO analysis revealed that the cellular targets clustered into 159 biological processes, including those involved in cellular composition (n=24) and molecular functions (n=42, P<0.01). KEGG pathway analysis revealed 85 (P < 0.01) pathways related to cancer. The active compounds in rhubarb target TP53, AKT1 and PIK3CA. Rhubarb therefore regulates cancer development through an array of biological pathways.

## Introduction

Cancer is a systemic disease that remains a major threat to human life[1]. The incidence of cancer continues to rise annually across the globe, with up to 23.91% of malignant tumors now thought to lead to cancer-related death[2]. The current front-line treatments for cancer include radiotherapy, surgery and chemotherapy[3]. Despite advances, many anti-cancers drugs lack efficacy and cause serious adverse reactions[4]. Traditional Chinese medicine has emerged as an alternative anti-cancer strategy due to its ability to prevent tumor cell DNA synthesis, promote apoptosis, induce tumor cell differentiation, inhibit tumor cell proliferation and metastasis, and inhibit angiogenesis[5–6].

Rhubarb originates from the dry root and rhizome of Rheum tanguticum Maxim. Ex BALF. Rheum officinale Baill or Rheum palmatum L.In TCM, rhubarb has been shown to promote detoxification, blood homeostasis, reduce diarrhea and prevent solid tumor growth in the abdomen. For example "Shennong materia medica classic" recorded that: "rhubarb tastes bitter, has a cold nature, and can be used to maintain blood homeostasis, and prevent addiction and accumulation”[7–8]. The main chemical constituents of rhubarb include anthracene derivatives, stilbenes, and tannins[9]. Amongst these components, emodin promotes hepatoma cell apoptosis and suppresses cell growth[10], whilst Aloe emodin (AE) inhibits the proliferation of an array of tumor cells[11]. Despite this knowledge, the molecular mechanisms governing the anticancer activity of rhubarb remain largely undefined.

Network pharmacology uses a systems biology approach to analyze specific signal nodes within a biological system to identify the cellular targets of drug therapy [12–13]. In this study, the key targets of rhubarb were investigated via a PPI network and "rhubarb drug component target" network model. Identified targets were analyzed using GO and KEGG databases to systematically define the material basis for the anti-cancer activity of rhubarb.

## Materials and methods

### Screening of the active components of rhubarb

The TCMSP database was used to investigate the relationship between Chinese herbal medicine, the targets of its components, and disease processes. Each component was evaluated for human absorption, distribution, metabolism and excretion. To avoid omission of the reported chemical components of rhubarb, oral bioavailability of the drug components were set to (OB≥20%), and drug similarities were set to (DL≥0.1) [14–15]. The chemical structure of rhubarb was retrieved using the PubChem platform[16].Swiss target predictions were used to identify cellular targets[17].

### Assessment of disease targets

TTD and OMIM databases were searched for “cancer” and “neoplast” to identify the genes related to tumor development[18–19]. Retrieved data were merged, duplicates were removed, and the intersection of the chemical targets was obtained using the Funrich platform[20]. Venn diagrams were constructed to reveal the key anticancer targets of rhubarb.

### Construction of the drug-disease database

Cytoscape3.6.1 was used to define the active components and anticancer targets of rhubarb. Nodes in the network represent rhubarb components, active ingredients and key target genes.

### Construction of the protein-protein interaction network

The String database was used to analyze the key cellular targets of rhubarb. Cytoscape3.6.1 was used to construct the PPI network. Node colors represent the importance of each target.

### Bioinformatics

The David database was used for GO and KEGG pathway analysis (P < 0.01). GO analysis included three modules: biological processes, molecular functions and cell composition as key targets for rhubarb. Prism software and Omicshare software were used for data analysis.

## Results

### Chemical components of rhubarb

A total of 92 chemical components of rhubarb were obtained. According to the set conditions (OB≥20% and DL≥0.1), 33 active substances were identified and numbered, including anthraquinone, flavonoids and tannins(Table 1).

**Table 1.** The detailed information about candidate compounds of Rheum.

| NO | Mol ID | Molecule Name | OB(%) | DL |
| --- | --- | --- | --- | --- |
| 1 | MOL002230 | (+)-Catechin-pentaacetate | 27.58 | 0.77 |
| 2 | MOL002235 | EUPATIN | 50.8 | 0.41 |
| 3 | MOL002239 | 5-acetyl-7-hydroxy-2-methyl-chromone | 30.25 | 0.1 |
| 4 | MOL002240 | 5-Carboxy-7-hydroxy-2-methyl-benzopyran-gamma-one | 34.4 | 0.11 |
| 5 | MOL002243 | Anthraglycoside B | 27.06 | 0.8 |
| 6 | MOL002244 | Chrysophanol glucoside | 20.06 | 0.76 |
| 7 | MOL003353 | Emodin anthrone | 24.72 | 0.21 |
| 8 | MOL002248 | Gallic acid-4-O-(6'-O-galloyl)-glucoside | 27.06 | 0.67 |
| 9 | MOL002251 | Mutatochrome | 48.64 | 0.61 |
| 10 | MOL002258 | Physcion-9-O-beta-D-glucopyranoside_qt | 20.3 | 0.3 |
| 11 | MOL002259 | Physciodiglucoside | 41.65 | 0.63 |
| 12 | MOL002260 | Procyanidin B-5,3'-O-gallate | 31.99 | 0.32 |
| 13 | MOL002262 | 5-[(Z)-2-(3-hydroxy-4-methoxy-phenyl)vinyl]resorcinol | 76.25 | 0.15 |
| 14 | MOL002268 | Rhein | 47.07 | 0.28 |
| 15 | MOL002270 | Rheinoside A_qt | 26.28 | 0.75 |
| 16 | MOL002274 | (9S)-9-[(9R)-2-carboxy-4,5-dihydroxy-10-oxo-9H-anthracen-9-yl]-4,5-dihydroxy-10-oxo-9H-anthracene-2-carboxylic acid | 27.75 | 0.57 |
| 17 | MOL002276 | Sennoside E_qt | 50.69 | 0.61 |
| 18 | MOL002280 | Torachrysone-8-O-beta-D-(6'-oxayl)-glucoside | 43.02 | 0.74 |
| 19 | MOL002281 | Toralactone | 46.46 | 0.24 |
| 20 | MOL002284 | PIT | 72.29 | 0.13 |
| 21 | MOL002288 | Emodin-1-O-beta-D-glucopyranoside | 44.81 | 0.8 |
| 22 | MOL002293 | Sennoside D_qt | 61.06 | 0.61 |
| 23 | MOL002294 | Strumaroside | 20.78 | 0.67 |
| 24 | MOL002296 | Daucosterol | 20.18 | 0.69 |
| 25 | MOL002297 | Daucosterol_qt | 35.89 | 0.7 |
| 26 | MOL002298 | Aloeemodin | 20.65 | 0.24 |
| 27 | MOL002303 | palmidin A | 32.45 | 0.65 |
| 28 | MOL000358 | beta-sitosterol | 36.91 | 0.75 |
| 29 | MOL000471 | aloe-emodin | 83.38 | 0.24 |
| 30 | MOL000472 | Emodin | 24.4 | 0.24 |
| 31 | MOL000476 | Physcion | 22.29 | 0.27 |
| 32 | MOL000554 | gallic acid-3-O-(6'-O-galloyl)-glucoside | 30.25 | 0.67 |
| 33 | MOL000096 | (-)-catechin | 49.68 | 0.24 |

### Screening of cellular targets

The target genes of the 33 active substances were screened in the Swiss Target Prediction database, revealing 398 genes using Homo sapiens as the search term. A total of 1018 tumor targets were then screened using the TTD and OMIM platform. Target genes were matched using the Funrich platform and Venn maps were constructed. These analyses 116 genes that were related to the targets of cancer treatment, including Aurka (Aurora kinase A-Interacting protein) (m230), HDAC6 (histone deacetylase 6) (m231), and VEGFR1 (vascular endothelial growth factor receptor 1(232)(Fig 1).

**Figure 1.**
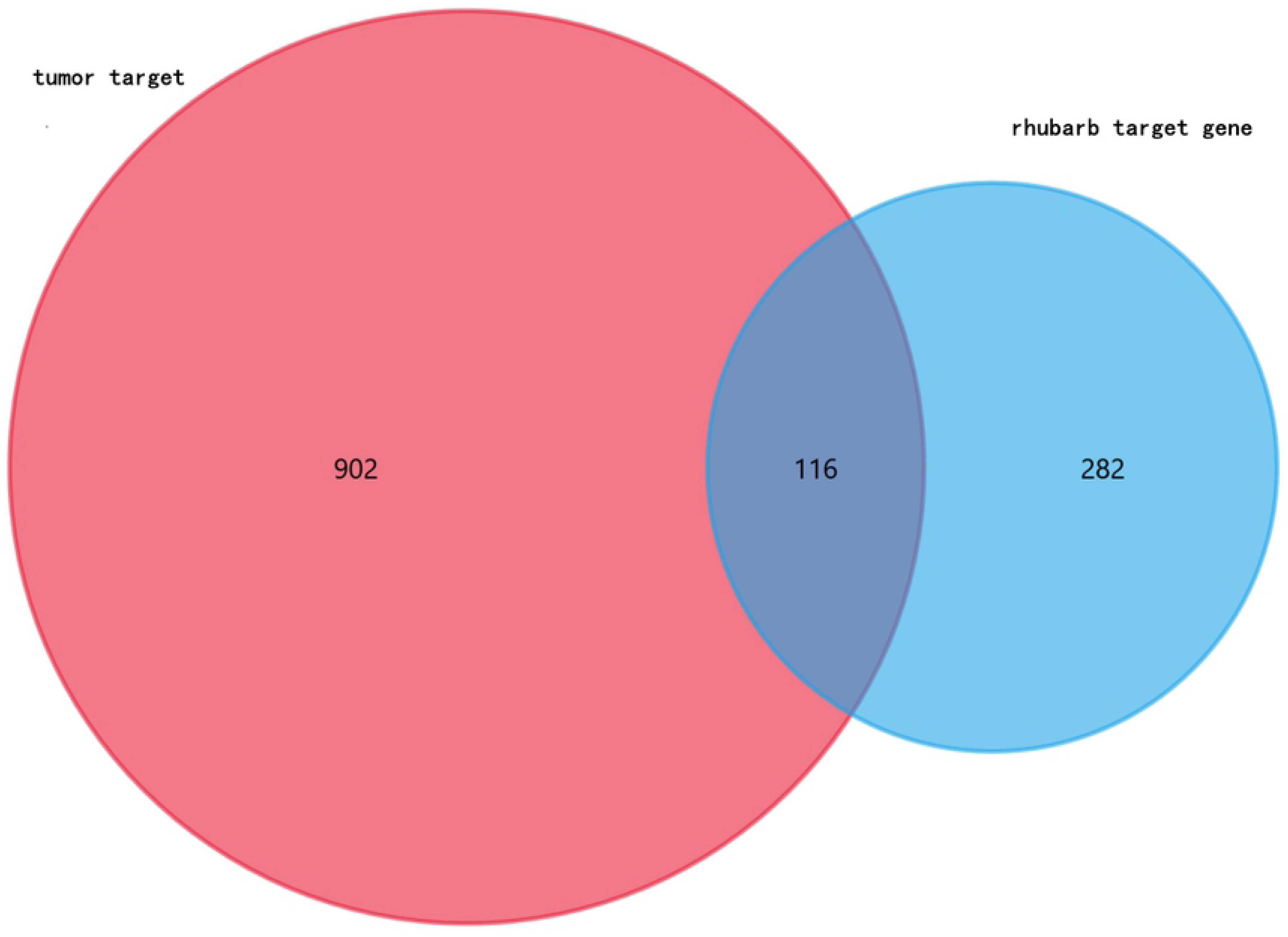

### PPI network

The PPI network included 103 nodes, including 1 medicinal material node, 17 compound nodes and 85 target nodes. The top five compounds were emodin anthrone, eupatin, physcion, catechin pentaacetate and palmidin with occurrences of 25, 29, 21, 16 and 36, respectively. Analysis the perspective of targets revealed Top1, Adora3, Met, Adora2b and EGFR as the top 5 targets, with degree values of 7, 7, 6, 6 and 6(Fig 2).

**Figure 2.**
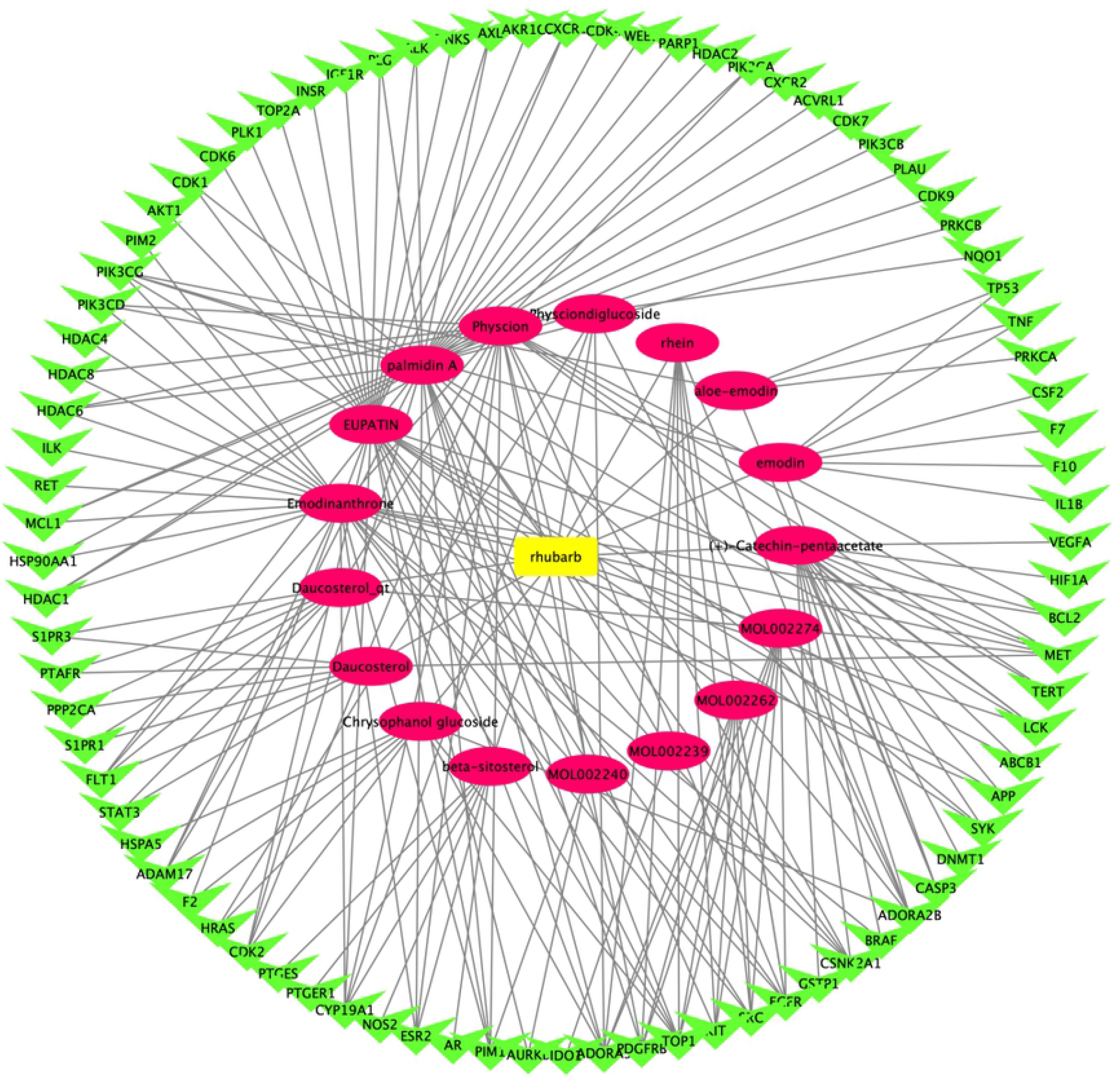

### Network modules

Key genes were imported in the String database for analysis. The network results showed revealed 101 nodes (action targets), with an average connectivity value of 10.752. The color of each node represents the importance of the target. The top 10 target genes were TP53, AKT1, STAT3, PIK3CA, HRAS, VEGFA, SRC, hsp90aa1, EGFR, pik3cb, with degree values of 46, 40, 39, 37, 36, 33, 33, 32, 29 and 22, respectively(Fig 3).

**Figure 3.**
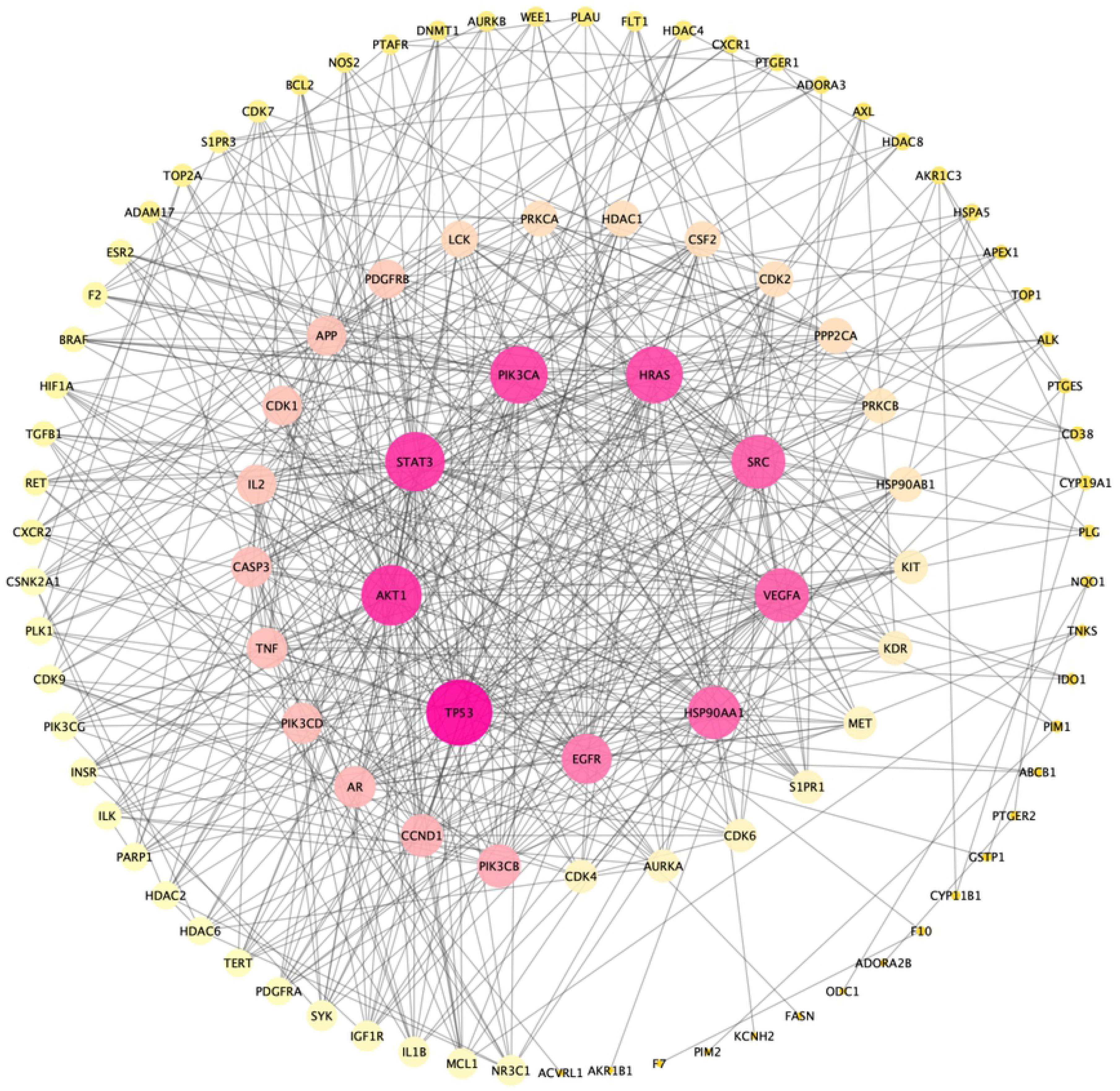

### Go and KEGG pathway analysis of the target genes

Go analysis showed that the target genes were enriched in 159 biological processes (BP) including the positive regulation of cell migration, protein phosphorylation and cell proliferation(Fig 4); 24 were enriched in cell composition (CC), including the plasma membrane, cytosol, nucleoplasm (Fig 5); and 42 were enriched in molecular functions (MF)(Fig 6); including ATP binding, protein binding, and enzyme binding. KEGG pathway analysis showed that the target genes were enriched in 89 cancer-related pathways, including Pathways in cancer, endometrial cancer, Small cell lung cancer, Non-small cell lung cancer, pancreatic cancer, Colorectal cancer, Bladder cancer, Prostate cancer, and Thyroid cancer(Fig 7).

**Figure 4.**
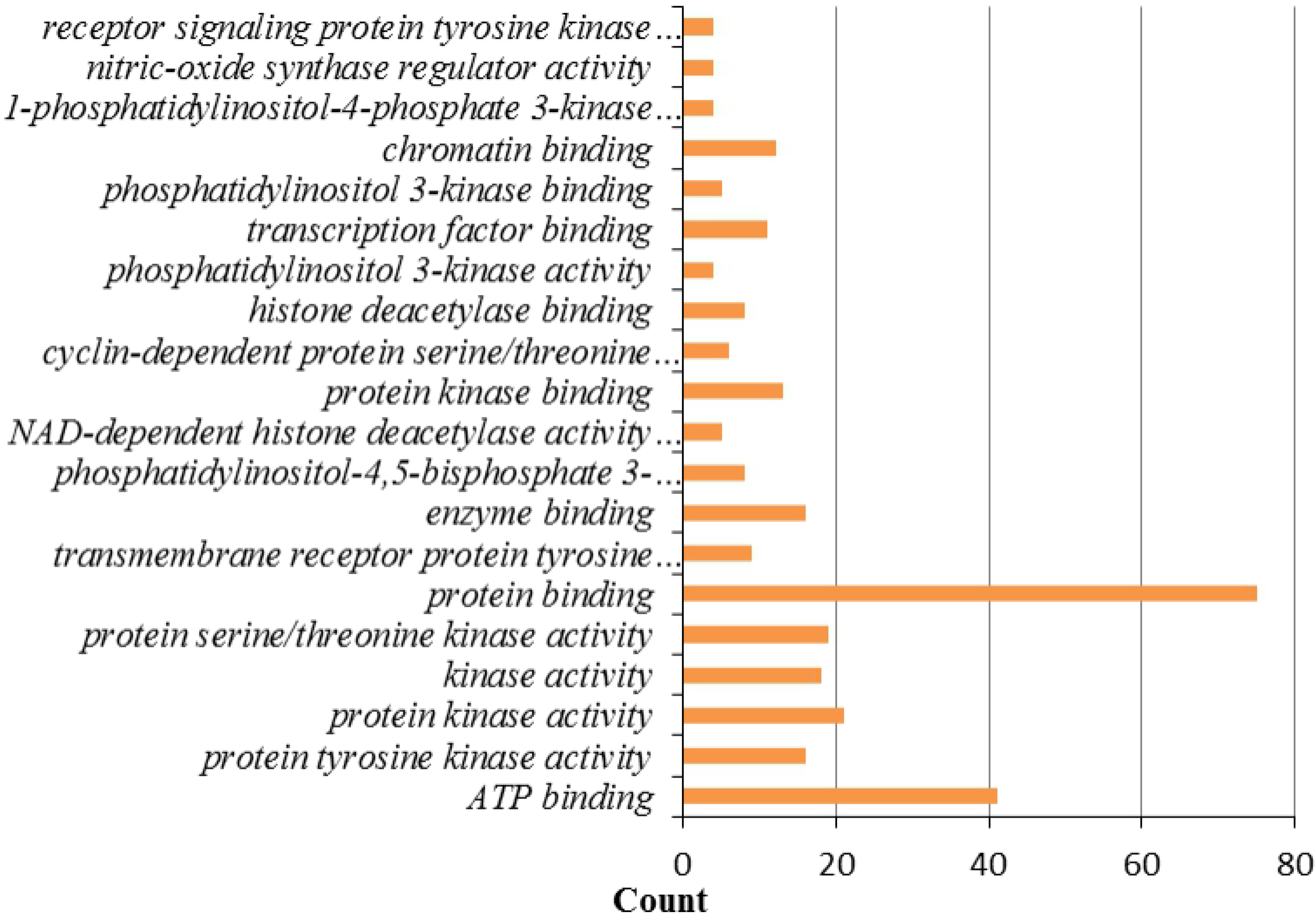

**Figure 5.**
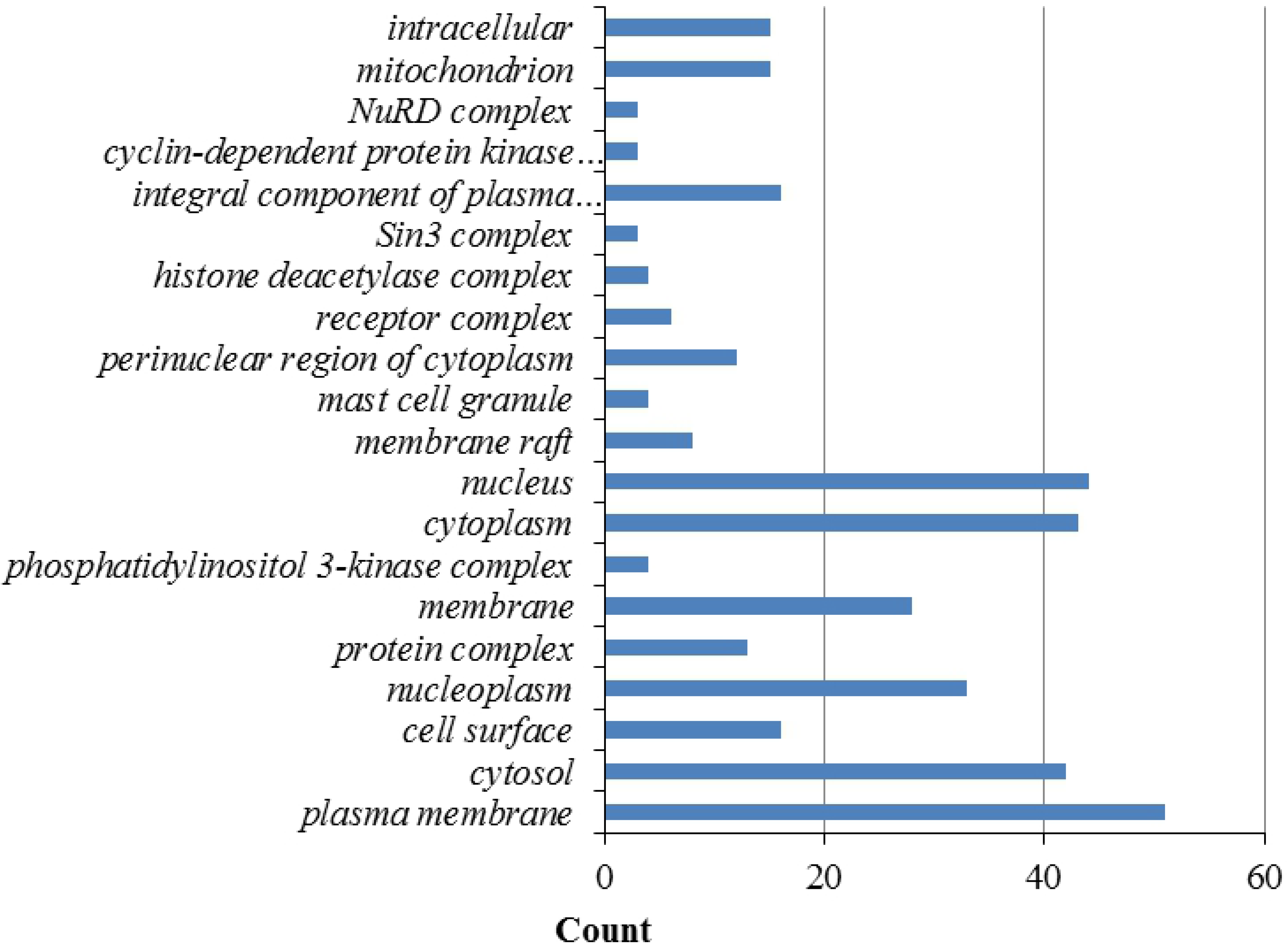

**Figure 6.**
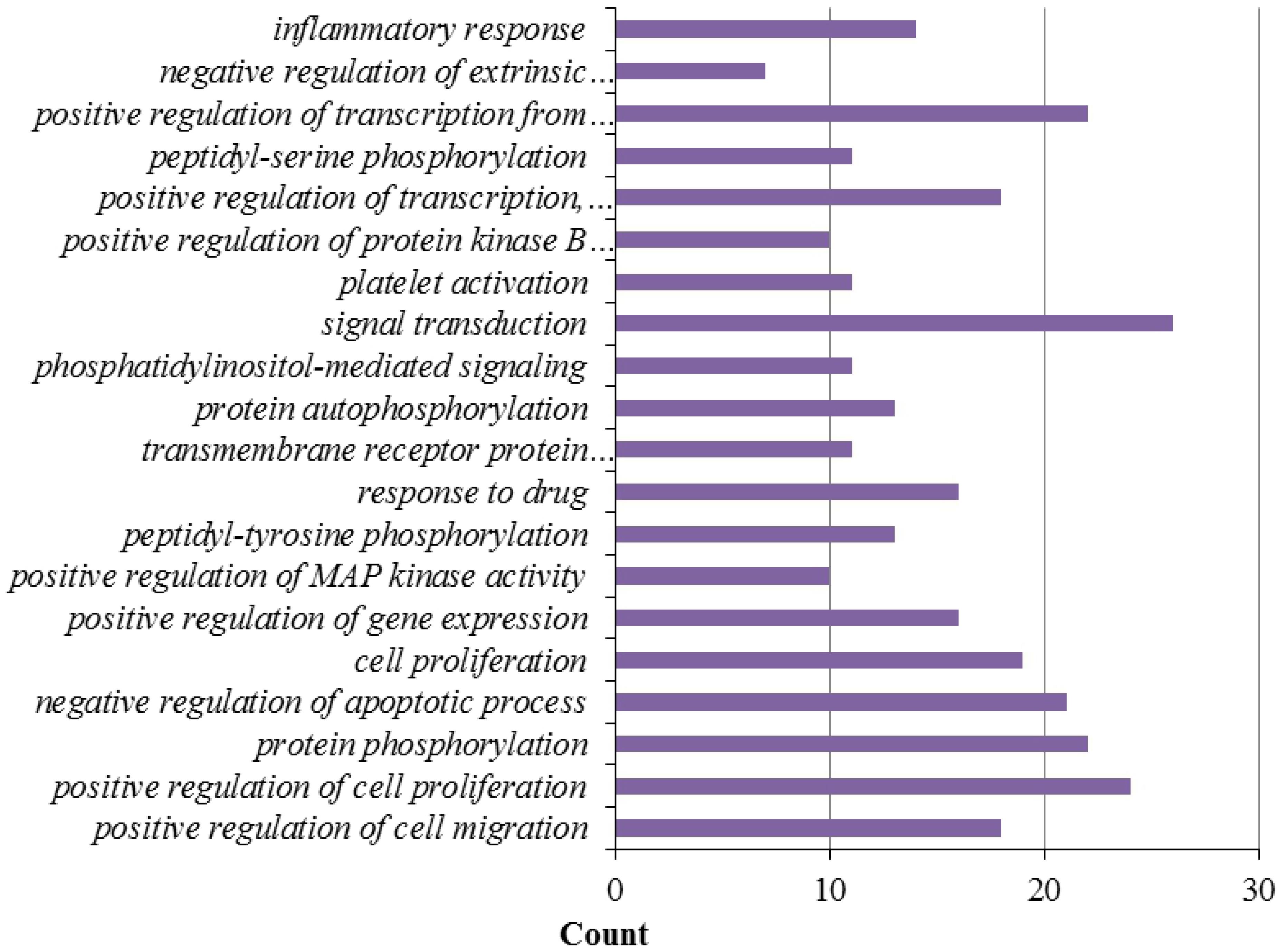

**Figure 7.**
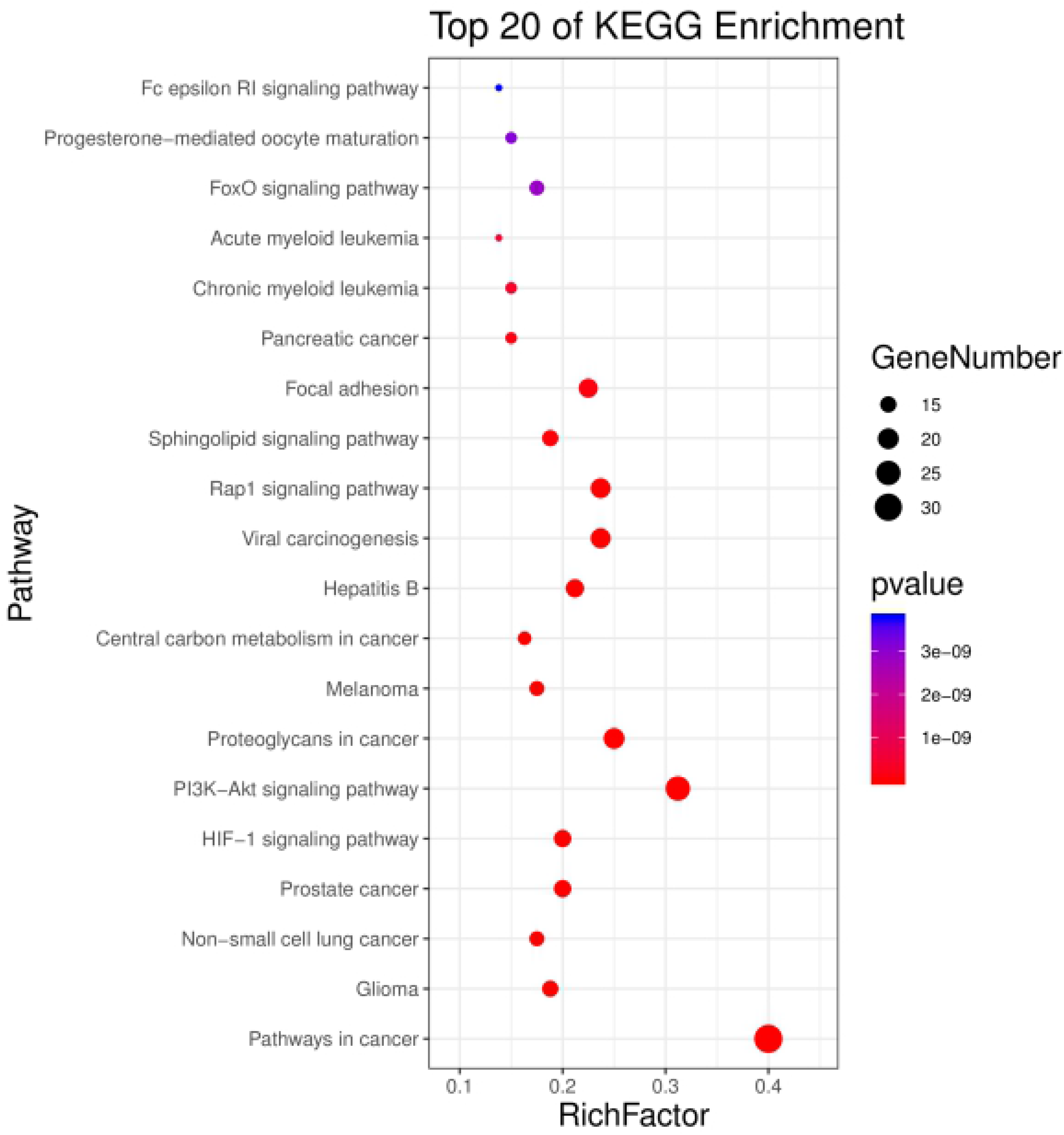

## Discussion

In this study, we show that anthraquinones, stilbenes and tannins form the active anticancer components of rhubarb. Emodin can suppress the growth of pancreatic cancer cells and reduce the expression of macrophage migration inhibitory factor, thereby promoting tumor cell apoptosis. Aloe emodin can inhibit the growth of mouse cervical cancer U14 cell transplantation tumors[21–22], inhibit the proliferation of CNE2 cells, and suppress the growth of the human malignant melanoma cell line A375[23]. β-sitosterol can inhibit the proliferation and differentiation of H22 transplanted tumors[24–25]. Eodin methyl inhibits the growth of colorectal and pancreatic cancer cells[26]. Emodin methyl ether can reduce the survival rates of pancreatic and breast cancer cells by suppressing the expression of cyclin D1, cyclin A, CDK4, CDK2 and c-myc[27].Rhein also prevents breast cancer development[28]. Analysis of the datasets revealed Top1, adora3, met, adora2b and EGFR as the five key targets of rhubarb. Met regulates the proliferation, survival, angiogenesis, invasion and metastasis of a variety of tumors. Abnormal Met expression promotes drug resistance and treatment failure[29]. EGFR activation regulates MAPK and PI3K/Akt signaling, promoting cell proliferation and differentiation. The mutation and abnormal expression of EGFR is related to the progression of non-small cell lung cancer (NSCLC), head and neck tumors, and genitourinary cancer[30]. These results suggest that rhubarb exerts its anti-tumor effects through multiple-components and cellular targets.

We identified 116 cellular targets of rhubarb through mapping the targets of the active ingredients with known tumorigenic host factors. Further analysis of the protein-protein interaction networks of the target genes showed that TP53, AKT1, STAT3, PIK3CA, HRAS, VEGFA, SRC, hsp90aa1 and EGFR play an important role in the network. Amongst the targets, the most frequent mutations in human tumor tissue occurred in TP53. Wild type *p53* is a sequence specific transcription factor that is activated in response to DNA damage, carcinogenic signal transduction, and nutrient depletion. Active *p53* promotes cell senescence, cell death, aging, and metabolic changes[31].Mutations in *p53* lead to a loss of tumor suppression, leading to oncogene activity, cell proliferation, invasion, metastasis and chemotherapy resistance, most notably in epithelial ovarian tumors[32].Mutant *p53* promotes the malignant progression of ovarian cancer by regulating the expression of tumor promoting genes such as TMEFF1[33–34]. In recent years, cell stimulating and regulatory factors have been shown to inhibit malignant progression by inhibiting the MDM2-p53 feedback loop. The ribosomal proteins RPS- MDM2 are a newly discovered axis that influence *p53* activity[35]. RPS inhibits the E3 ubiquitinase activity of MDM2, thereby stabilizing *p53*[36].To-date, 16 ribosomal proteins (RPS) have been shown to interact with MDM2 to activate *p53*. Amongst them, rpl5 and rpl11 are the most important sensors and effectors of ribosome stimulation. The Rps-mdm2-p53 axis therefore effectively prevents uncontrolled cell growth due to abnormal carcinogenic activity[37–38].

The human *EGFR* gene is located in the 7p12-14 region of the short arm of chromosome 7, consisting of 28 exons. EGFR is a transmembrane protein composed of 1210 amino acids. The extracellular domain mediates ligand-binding which activates the intracellular kinase domain[39].*EGFR* mutations are frequently observed in Asian women with adenocarcinoma who lack a history of smoking[40–41]. The most common mutations are the deletion of exon 19 (del19) and a point mutation within exon 21 (L858R) which accounts for ~90% of all polymorphisms[42].When EGFR binds to the epidermal growth factor (EGF), receptor dimerization results in the activation of the tyrosine kinase domain[43], and phosphorylated tyrosine residues recruit downstream adaptor proteins to activate specific signaling pathways, including PI3K-Akt, Ras-Raf-MEPK-ERK, and STAT family members, resulting in cell growth and differentiation. The inhibition of angiogenesis can suppress the growth, invasion, and metastasis of lung cancer cells, particularly NSCLC[44]. An array of cancer cells overexpress EGFR, highlighting its importance as an anticancer target. Advanced NSCLC patients with EGFR/TP53 co-mutations also respond poorly to TKI therapy[45–47].

The PI3K/Akt axis regulates malignant proliferation, metastasis, angiogenesis and chemoradiotherapy. This pathway represents one of the three major cell signaling axis that regulate IL-6 and its biological functions. PI3K/Akt signaling prevents the initiation of programmed cell death, promotes tumor cell invasion, and induces the production of HIF-1α and VEGF, thereby promoting tumor angiogenesis[48–50].

HRAS is one of the three members of ras gene family[51]. The mutation and activation of *Ras* weakens its ability to bind to guanosine diphosphate (GDP). Mutated Ras binds guanosine triphosphate (GTP) and self-activates in the absence of external growth signals. GTPase activity can reduce *Ras* activity by enhancing the dissociation of Ras and GTP. Continuous Ras activation promotes PLC activity, leading to uncontrolled cell proliferation[52]. The carcinogenic effects of *HRAS* manifest through protein modifications and enhanced expression[53–55]. The HRAS T81C polymorphism does not alter the amino acid sequence of Ras, but induces its overexpression, thereby influencing cancer susceptibility[56].

Tumor cells directly or indirectly recruit host cells and secrete a variety of angiogenesis promoting factors, amongst which vascular endothelial growth factor (VEGF) is a key mediator[57].The relative molecular weight of VEGF is ~34-45KD with seven related gene families described, including VEGFA, VEGFB, VEGFC, VEGFC, VEGVEE, VEGGF and placental growth factor (PIGF). Amongst them, VEGFA is most abundant in tissues and cells and plays an important role in angiogenesis. VEGFA (sometimes referred to as BEGF)[58], can bind to the vascular endothelial cell receptor (VEGFR), to induce the proliferation, survival and migration of endothelial cells, thereby mediating angiogenesis[59]. It has been reported that Berna is closely related to the growth and metastasis of cancers[60].

SRC is a ~60kD phosphorylated protein encoded by the proto-oncogene *c-Src*, which is the first cancer protein with tyrosine protein kinase activity[61].As a non-receptor tyrosine kinase, SRC widely participates in tumorigenesis and development through the regulation of tumor cell proliferation, differentiation, migration, invasion and angiogenesis[62].Activated SRC stimulates STAT3 through a JAK-independent pathway in hepatoma cells[63]. Activated SRC also enhances the transcription of hepatocyte growth factor gene. SRC induces STAT3 phosphorylation to enhance its affinity to DNA, synergistically enhancing the activity of the HCF promoter to enhance HCF transcription[64–65]. Rhubarb inhibits STAT3 signaling both *in vitro* and *in vivo*, inhibiting cancer cell proliferation, migration, invasion, and angiogenesis[66].

Heat shock protein 90 (Hsp90) is a highly conserved protein chaperone in eukaryotes[67–68]. A variety of Hsp90 subtypes exist, including Hsp90α and Hsp90β, endoplasmic reticulum GRP94 protein, and the mitochondrial protein TRAP1[69]. Hsp90αis encoded by the *hsp90aa1* gene. Human *hsp90aa1* is encoded by a complementary chain on chromosome 14q32.33. As a tumor promoter, Hsp90 interacts with a variety of oncogenic proteins and participates in malignant transformation and tumor development[70–71]. Up to 115 mutations in the open reading frame of *hsp90aa1* have been identified through next-generation-sequencing of tumor tissue and cell lines. However, the effects of these mutations on the function of *hsp90aa1* require clarification. Homozygous deletions of the *hsp90aa1* gene in tumor tissue lead to a lower occurrence of malignancy. Clinical studies have shown the association of *hsp90aa1* with a poor prognosis in 206 patients with gastric cancer[72]. Conversely, Hsp90aa1 deletions can be used as a biomarker for a favorable prognosis in biopsy specimens of tumor tissue[73–74]. The biological functions of Hsp90α and Hsp90β differ. Unlike Hsp90β, Hsp90α plays a role in wound repair and inflammation in tumor cells, influencing their migratory and extravasation phenotypes[75]. Recent studies on prostate cancer have shown that extracellular Hsp90α promotes chronic inflammation in tumor associated fibroblasts. Changes in the extracellular environment of malignant tumors can also promote the malignant progression of prostate cancer[76].Extracellular Hsp90α induces inflammation through the activation of NF-kB and STAT3 transcription. NF-KB also induces the expression of Hsp90α[77]. Exocrine Hsp90α stimulates fibroblasts to secrete intracellular Hsp90α through an autocrine and paracrine feedback loop, leading to the generation of an inflammatory storm around malignant tumor cells. These features may explain the correlation between Hsp90α expression and malignancy, which requires investigation in future studies[78].

We found that TP53, AKT1, STAT3, PIK3CA, HRAS, VEGFA, SRC, hsp90aa1 and EGFR were related to the malignant behavior of tumor cells in both stressed and non-stressed states. We systematically analyzed the antitumor effects of rhubarb, and identified key roles for emodin, aloe emodin, β-sitosterol, and emodin methyl ether in NSCLC, prostate cancer, pancreatic cancer, and endometrial cancer. We further identified TP53, AKT1, HRAS and EGFR as the cellular targets of these rhubarb constituents. Some limitations should be noted including incomplete data collection. These findings do however provide a basis for future studies on the anticancer activity of rhubarb.

## Supporting information

**S1 Fig. Matching of target genes between disease and *rhubarb*. (TIF)**

**S2 Fig. Drug-component-target network of active ingredients of *rhubarb*. (TIF)**

**S3 Fig. Protein interaction network of *rhubarb*.(TIF)**

**S4 Fig. Molecular functions analysis of potential targets in *rhubarb*. (TIF)**

**S5 Fig. Cell components analysis of potential targets in *rhubarb*. (TIF)**

***S6 Fig*. Biological process analysis of potential targets in *rhubarb*. (TIF)**

**S7 Fig. KEGG pathway analysis of potential targets in *rhubarb*. (TIF)**

## Acknowledgments

None

## Author contributions

Conceptualization: Hu Junrui, Duan yongqiang and Xi shanshan.

Data curation: Huang Rui and Cui Gongning.

Funding acquisition: Duan Yongqiang and Wu Hongyan.

Methodology: Ma Jun and Bai Min.

Project administration: Duan Yongqiang and Wu Hongyan.

Software: Luo Qiang, Xi shanshan. and Hu Junrui

Validation: Duan Yongqiang

Writing – original draft: Hu Junrui, Duan yongqiang and Xi shanshan.

Writing – review & editing: Cui Gongning and Wu Hongyan

